# A Novel Activation Maximization-based Approach for Insight into Electrophysiology Classifiers

**DOI:** 10.1101/2021.10.10.463830

**Authors:** Charles A. Ellis, Mohammad S.E. Sendi, Robyn Miller, Vince Calhoun

## Abstract

Spectral analysis remains a hallmark approach for gaining insight into electrophysiology modalities like electroencephalography (EEG). As the field of deep learning has progressed, more studies have begun to train deep learning classifiers on raw EEG data, which presents unique problems for explainability. A growing number of studies have presented explainability approaches that provide insight into the spectral features learned by deep learning classifiers. However, existing approaches only attribute importance to different frequency bands. Most of the methods cannot provide insight into the actual spectral values or the relationship between spectral features that models have learned. Here, we present a novel adaptation of activation maximization for electrophysiology time-series that generates samples that indicate the features learned by classifiers by optimizing their spectral content. We evaluate our approach within the context of EEG sleep stage classification with a convolutional neural network, and we find that our approach is able to identify spectral patterns known to be associated with each sleep stage. We also find surprising results suggesting that our classifier may have prioritized the use of eye and motion artifact when identifying Awake samples. Our approach is the first adaptation of activation maximization to the domain of raw electrophysiology classification. Additionally, our approach has implications for explaining any classifier trained on highly dynamic, long time-series.

## I. Introduction

Electroencephalography (EEG) is a popular modality for exploring brain function that has been applied within the context of many neurological and neuropsychiatric disorders [1]–[3]. A key characteristic of EEG data is its rich frequency-domain. As the field of deep learning has evolved, deep learning methods have been increasingly applied to EEG time-series [4]–[7], and in response, a growing group of studies have presented explainability approaches specifically adapted to providing insight into the spectral features learned by neural networks. In this study, we present a novel Frequency-based Activation Maximization Explainability (FAME) approach that provides in-depth insight into the spectral features learned by deep learning classifiers and that generates highly realistic time-series signals with features demonstrating the characteristic patterns learned by deep learning classifiers.

Machine learning methods have now been used to gain insight into EEG spectra for the better part of two decades. Particularly before the rise of deep learning, it was more common to apply traditional interpretable machine learning approaches for insight into EEG spectra and the spectra of related modalities [8]. Machine learning classifiers were trained directly on EEG spectra, and a standard explainability approach could be applied. As deep learning has developed, many studies have applied deep learning approaches and deep learning explainability methods to analyze EEG spectra [1], [2]. These studies have a key shortcoming. They involve manual feature extraction and an inherent assumption that spectral features are the most worthwhile features that can be extracted. While spectral features can be very informative, other EEG waveforms can be useful in developing a high-performing classifier [9].

As a result, many studies have trained neural networks on raw EEG time-series. While the use of raw data can provide a richer feature space, the use of raw data also presents a unique challenge for explainability. Standard deep learning explainability approaches often do not provide useful insight into the features learned by classifiers for time-series. They either cannot be directly adapted or can only output importance values for each time point, which can be hard to understand. Because of this, there are very few studies that apply traditional deep learning explainability methods to EEG time-series [10]. As such, a growing group of studies have sought to develop methods that give insight into the spectral features learned by deep learning classifiers from raw time-series. With the exception of a study that visualized model weights [11], most of these methods have involved perturbation of the frequency domain and a subsequent examination of the effect upon the performance of the classifier. Those methods are collectively considered “feature attribution” methods [12]–[15]. Both global [13][15] and local approaches [16][17] have been developed. However, these methods only provide an estimate of importance. They do not indicate the spectral values learned by the classifier for each class, and furthermore, they have not provided insight into the relationships between frequencies that characterized the concept learned by the model for each class.

Prototyping methods could offer insight into the spectral values that the model associates with each class and that could provide insight into the relationships between frequency bands learned by the classifier. Prototyping methods can construct samples that fit the representation learned by the classifier for each class [18]. Activation maximization is a common prototyping approach that has mainly been used within the context of image analysis [19]. It has since been applied to time-series analysis in a small number of instances [4], [20], [21]. In two of these instances, it was only used for very short time-series (i.e., approximately 30 points in length) within the domain of activity classification [20], [21]. In the remaining instance, the authors input sinusoids into a model and examined the activation of the intermediate nodes of the model [4]. This provided insight into the frequencies identified by the model. However, because it required outputting activations for many nodes, the results were complicated and difficult to interpret. Additionally, they did not perform activation maximization for the final output nodes. In summary, although activation maximization has been applied in the time domain, it has either not been applied to long, spectrally rich time-series like those found in electrophysiology, or it has not been applied to the output layers of a model.

In this study, we develop FAME, a novel adaptation of activation maximization that maximizes the output of the final output layer and generates realistic samples for lengthy time-series. Moreover, Our approach provides insight into the key spectral patterns learned by a classifier for each class. our approach involves iteratively converting samples to and from the frequency domain, examining the gradient associated with the real and imaginary coefficients of each frequency, and updating the values based upon that gradient. We demonstrate the viability of our approach within the context of automated sleep stage classification with samples that have lengths of around 3,000 points. Importantly, sleep stages have well-characterized frequency signatures, making sleep stage classification an ideal opportunity for determining whether our approach uncovers realistic representations. After generating activation maximization samples, we compare our resulting samples to the training data and established sleep staging guidelines. We then apply a local perturbation-based spectral explainability approach for insight into the importance of the spectral features learned by our approach.

## II. Methods

In this section, we discuss our dataset, classifier, and novel explainability approach.

### A. Description of Data

In this study, we use Sleep Cassette data from the popular Sleep-EDF Database Expanded [22] that is publicly available on Physionet [23]. The dataset consists of 153 approximately 20-hour recordings collected from 78 participants. There are 41 female and 37 male participants. The participants have ages with a mean of 58.79 years and a standard deviation of 22.15 years. The data is recorded at a sampling rate of 100 Hertz. The dataset consists of two EEG electrodes. We use the FPz-Cz electrode because it has been shown to be sensitive to sleep staging in previous studies [2][24]. The data was annotated by domain experts in 30-second segments. We used Awake, Rapid Eye Movement (REM), non-REM 1 (NREM1), non-REM 2 (NREM2), non-REM 3 (NREM3), and non-REM 4 (NREM4).

### B. Description of Data Preprocessing

Using the expert-assigned labels, we separate the data into samples with a length of 30-seconds. To help alleviate class imbalances that occurred because the recordings covered a 20-hour period, we removed the *Awake* samples at the start of each recording. We further reduce the size of the Awake class by removing Awake samples from the end of the recording such that the *Awake* class and NREM2 class had a comparable number of samples. In accordance with the recommendations of the American Society for Sleep Medicine, we combine the NREM3 and NREM4 classes into a single NREM3 class [25]. After sample removal, we z-score the samples remaining from each recording. Our dataset consists of 215,668 samples with 39.43%, 9.98%, 32.05%, 6.05%, and 22.98% of the dataset being Awake, NREM1, NREM2, NREM3, and REM, respectively.

### C. Description of Classifier

We use a CNN classifier that was first developed in [26] and has since been used in a number of other studies [27], [28]. The architecture is shown in Figure 1. It is composed of 8 convolutional layers with intermixed max pooling, global max pooling, spatial dropout, and dropout. When training the classifier, we use 10-fold cross-validation in which 63, 7, and 8 participants are randomly assigned to the training, validation, and test groups, respectively. We use the Adam optimizer with an initial learning rate of 0.001 that decreases after every 5 consecutive epochs that occur without an increase in performance. We use class-weighted categorical cross-entropy loss to minimize the effects of class imbalance. When testing the classifier, we compute the F1 score, recall, and precision of each fold and subsequently compute their mean and standard deviation.

**Fig. 1.**
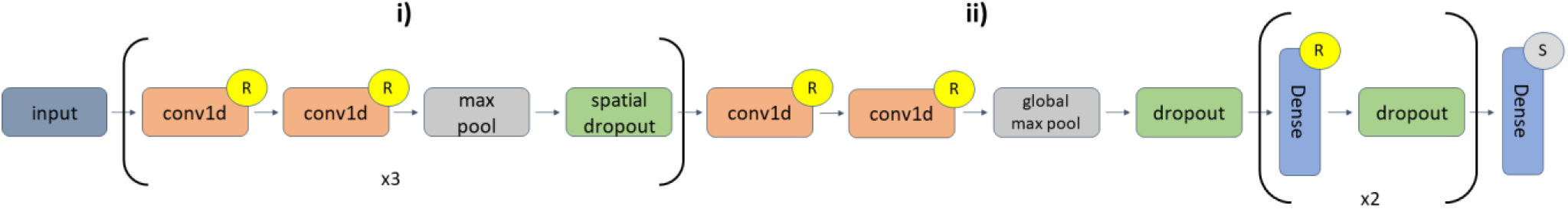
CNN Architecture. **i)** had 3 sets of 2 1-dimensional convolutional (conv1d) layers with ReLU activation functions followed by max pooling and spatial dropout. The conv1d layers of the first set had 16 filters and a kernel size of 5. The first max pooling layer had a pool size of 2, and the spatial dropout layer had a rate = 0.01. The conv1d layers of the second set had 32 filters and a kernel size of 3. The second max pooling layer had a pool size of 2, and the spatial dropout layer had a rate = 0.01. The conv1d layers of the third set had 32 filters and a kernel size of 3. The third max pooling layer had a pool size of 2, and the spatial dropout layer had a rate = 0.01. The two conv1d layers in **ii**) had 256 filters with a kernel size of 3. They were followed by global max pooling and dropout with a rate of 0.01. Afterwards were 2 sets of dense layers with 64 nodes each that were followed dropout with rates of 0.1 and 0.05 for the first and second dense layers, respectively. The last dense layer had 5 nodes. Centers with an “R” or “S” in the center indicate layers followed by ReLU or Softmax activation functions, respectively.

### D. Description of Explainability Approach

In this study, we develop a novel adaptation of activation maximization. FAME is particularly suited to long time-series. Our method can produce a sample that is representative of the features learned by the classifier for a particular class. It consists of a series of steps. (1) Our approach starts by initializing a sample for a particular class. The initialization involves converting all training samples for the respective class to the frequency domain via a Fast Fourier Transform (FFT), obtaining the median of the real and imaginary coefficients for each frequency, and converting the median back to the time domain. (2) The class-specific activation produced by the classifier for the sample is obtained. (3) From there, each of the real and imaginary values in the sample is perturbed by a value of 0.001, the resulting perturbed samples are fed to the classifier, and the change in activation for each perturbation is calculated. (4) After the change in activation is calculated, that can be combined with the original perturbation size to obtain a gradient. (5) We then find the direction of the gradient and modify the corresponding real or imaginary value by a predefined step size. (6) We repeat steps 2 through 6 for a number of predefined steps and select the sample with the corresponding largest activation. (7) We perform a grid search across multiple real and imaginary step size values and select the activation that is highest across all step sizes. We maintain boundary conditions throughout the optimization process. If samples obtain real or imaginary coefficients for a positive frequency that are outside the minimum or maximum coefficients for that class of training data, those coefficients and their corresponding negative coefficients are reset to the minimum or maximum class value.

In our analysis, we examine step sizes of 0.3, 0.9, 1.5, 2.1, 2.7, 3.5, and 4.1 for the real coefficients. We use step sizes of 0.003j, 0.009j, 0.09j, 0.5j, 1.1j, 1.7j, and 2.1j for the imaginary coefficients. We use 700 steps. When using our activation maximization approach, we use the training data from the cross-validation fold with the highest weighted F1-score on the test data to obtain values for initialization and boundary conditions. After using activation maximization to generate samples for each class, we use a local frequency perturbation technique that we developed in another study to identify the relative importance of each of the canonical frequency bands to the generated samples [16]. We use 5 frequency bands: δ (0 – 4 Hz), θ (4 – 8 Hz), α (8 – 12 Hz), β (12 – 25 Hz), and γ (25 – 50 Hz). Before developing our novel frequency-based activation maximization approach, we also attempted time-based activation maximization with an established library. However, given the poor results that we obtained for time domain-based activation maximization, we do not include them in this study.

## III. Results and Discussion

In this section, we describe and discuss our classification performance and the samples generated with our novel activation maximization approach. We describe how the generated samples relate to existing sleep literature in the time-domain and frequency-domain. We further identify and discuss the most important frequencies for the samples that we generate. We also elaborate upon the broader implications of our approach, its limitations, and next steps for extending it to other time-series problems. It should be noted that, unless otherwise stated, when we compare our activation maximization samples to existing literature, we are comparing them to the sleep scoring manual developed by the American Academy of Sleep Medicine [25]. This manual has also been referenced in a number of other sleep stage classification studies [9], [29], [30].

### A. Classification Performance

Table 1 shows the mean and standard deviation of the F1 score, recall, and precision of the classifier across all folds. It also shows the metrics for the fold with the top weighted F1-score that we used when performing activation maximization. The Awake class obtains the highest performance across metrics and for all folds and the top fold. This is appropriate given that the Awake class makes up a large portion of the dataset. Additionally, Awake periods tend to have EEG activity that is distinct from other sleep stages and can have artifacts from eye movement and other motion. NREM1 has the lowest performance across all metrics. This is partially attributable to NREM1 being one of the smallest classes in our dataset. Sleep stage classification studies have historically had difficulty classifying NREM1 [15], [31], [32], and some studies have specifically developed approaches with the goal of maximizing NREM1 performance [33]. After Awake, NREM2 has the highest F1 score and precision for the top fold and all folds. After Awake, NREM3 has the highest recall, and it has an F1 score and precision comparable to REM. The NREM3 class has the smallest number of samples of any class in the dataset. However, the classifier still performs well on the class, possibly because the waveform patterns associated with NREM3 are particularly distinct [25].

**TABLE I.**
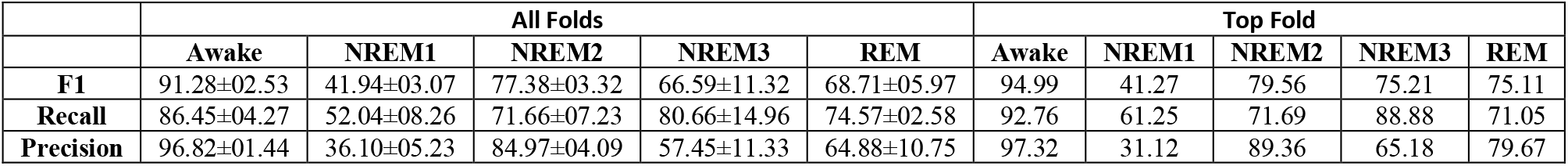
Classification Performance Results

|  | All Folds |  |  |  |  | Top Fold |  |  |  |  |
| --- | --- | --- | --- | --- | --- | --- | --- | --- | --- | --- |
|  | Awake | NREM1 | NREM2 | NREM3 | REM | Awake | NREM1 | NREM2 | NREM3 | REM |
| **F1** | 91.28±02.53 | 41.94±03.07 | 77.38±03.32 | 66.59±11.32 | 68.71±05.97 | 94.99 | 41.27 | 79.56 | 75.21 | 75.11 |
| **Recall** | 86.45±04.27 | 52.04±08.26 | 71.66±07.23 | 80.66±14.96 | 74.57±02.58 | 92.76 | 61.25 | 71.69 | 88.88 | 71.05 |
| **Precision** | 96.82±01.44 | 36.10±05.23 | 84.97±04.09 | 57.45±11.33 | 64.88±10.75 | 97.32 | 31.12 | 89.36 | 65.18 | 79.67 |

### B. Activation Maximization: Analysis of Activations

Fig. 2a visualizes the activations for each sample in the training dataset for their respective classes. Additionally, it shows the activations for each of the samples that we generate with our frequency-based activation maximization approach. These activations, their associated coefficient step sizes, and the number of steps required to reach peak activation are also shown in Table 2. Our Awake sample has an activation of 100%, with zero activation for other classes. It is reasonable that our approach yields an activation this high for Awake given our initialization approach and boundary conditions. The initialization and boundary conditions are based on the Awake samples in the training dataset, and the Awake samples generally have activations near 100%. Additionally, our model has very high classification performance for Awake, and our algorithm takes 537 steps to arrive at a maximal activation. Our NREM1 sample has an activation of 86.62% for NREM1, and the rest of its activation is attributed to Awake. Interestingly, the activation for our NREM1 sample is far above the median activation for the NREM1 training data, and activation maximization manages to obtain a very high activation considering that the performance of the model for the class is very low. It is possible that our algorithm is able to find a higher activation more easily because the starting activation at initialization is near the median activation for the class, which is supported by the number of steps required to reach maximal activation. The activation of the NREM2 sample is the lowest of all classes at 51.85% and is below the median activation for the NREM2 training samples. The sample produces strong activation for the NREM1 class and to a lesser degree for REM. However, it nonetheless activates more for NREM2 than it does for any other class. This level of activation is odd, given that the classifier performed better for NREM2 than many of the other classes. However, the classifier has higher or comparable levels of recall for NREM3 and REM than it did for NREM2. The relative low level of activation for NREM2 could be attributable to the starting activation, which is relatively low. The activation for our NREM3 sample is quite high at 86.82% with a small amount of activation for NREM2. Its activation is not far below the median activation for the NREM3 training data, and the high activation may also be explained by the high recall of the classifier for NREM3. Our REM sample obtains an activation of 69.72% for REM with some activation for NREM1. Its activation is slightly below the median activation for the REM training data, which could be attributable to its relatively low activation at initialization. In all cases, our approach learns spectral patterns associated with each class that obtain activations well above chance level.

**Fig. 2.**
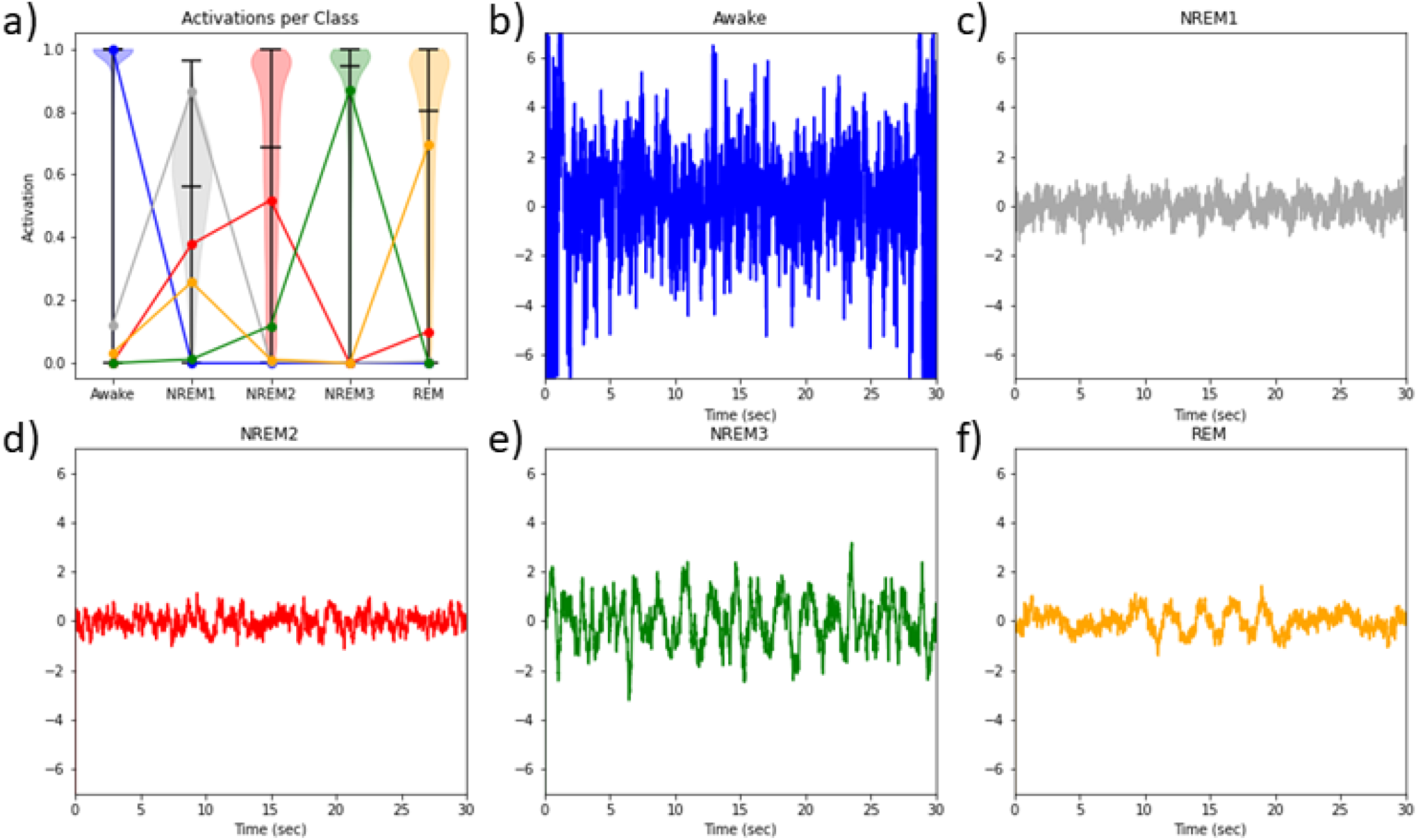
Sample Activations and Samples Generated with Activation Maximization. Panel a) shows a violin plot of all of the training samples within each class. It also shows the activations for each sample generated via activation maximization across all classes (i.e., colored lines and dots across classes). Panels b) through f) show the samples generated via activation maximization for each class. The y-axes are standardized for an easier comparison of the samples. There are minor edge effects for several of the classes that are difficult to see given the x-axes. It should be that blue, gray, red, green, and orange refer to Awake, NREM1, NREM2, NREM3, and REM, respectively.

**TABLE II.** Activation Maximization Results

|  | Awake | NREM1 | NREM2 | NREM3 | REM |
| --- | --- | --- | --- | --- | --- |
| **Starting Activation** | 02.11 | 56.42 | 39.08 | 74.44 | 38.50 |
| **Peak Activation** | 100.00 | 86.62 | 51.85 | 86.82 | 69.72 |
| **Number of Steps** | 537 | 18 | 54 | 13 | 124 |
| **Real Step Size** | 3.5 | 2.1 | 0.9 | 2.7 | 0.3 |
| **Imaginary Step Size** | 0.5j | 1.7j | 0.09j | 1.1j | 0.05j |

### C. Activation Maximization: Analysis of Time Domain

Fig. 2b through 2f visualize the time-series that resulted from our spectral activation maximization approach. In contrast to samples that could be generated with a traditional activation maximization approach, our samples generally look fairly realistic. NREM3 is typically characterized by δ activity, which has higher amplitude than other bands [25]. Similar to canonical sleep data, NREM3 had larger amplitudes and lower frequencies than NREM1, NREM2, and REM. Also, there seems to be a visible slowing in frequency from NREM1 to NREM2 to NREM3 as would be expected [25]. REM seems to have higher frequencies around the start and end of the sample and some lower frequency activity around the middle of the sample. This fits with the low amplitude, mixed frequency activity typically associated with REM. Although the Awake sample has the highest activation, it seems to be atypical. Awake periods typically have higher frequency, low amplitude EEG activity. Awake periods also typically have substantial amounts of eye movement that appears in frontal EEG electrodes unless the data is processed to remove them.

### D. Activation Maximization: Analysis of Frequency Domain

Fig. 3 shows the scaled power within each frequency band for the samples that we generated via activation maximization and for the training samples that we used when defining initialization values and boundaries conditions. Fig. 3a and Fig. 3b show the results for each of the activation maximization-based samples and for the training samples, respectively. To scale the spectral power values within each class into a comparable range, the frequency bands of each class are divided by the mean power of the δ-band of their respective classes. Note that for the most part the spectral power values of our samples generally follow the trend of exponential decay that is commonly found in biological signals and that is present in the samples shown in Fig. 3b. REM is the only class that seems to disrupt this pattern, as θ through γ bands have comparable levels of activity. As REM is often characterized as mixed frequency [25], that is not unsurprising. Awake and NREM1 both seems to have higher levels of γ activity than β activity, which departs from what is observed in the training sample distribution. Consistent with the training data, NREM3 seems to have a larger difference between δ and the other frequency bands than any of the other classes except Awake. This is also consistent with established literature which indicates that NREM3 is characterized by high levels of δ activity [25]. Relative to γ power, NREM1 also has larger θ power than the other classes, which is significant given that NREM1 is typically characterized by θ power [25]. Additionally, relative to δ, NREM2 has somewhat high levels of α and β activity, which are associated with the sleep spindles that appear in NREM2. NREM2 also has δ power that is moderately higher relative to the other frequency bands. Given that the k-complexes that appear in NREM2 occur in the δ band, that is not unexpected.

**Fig. 3.**
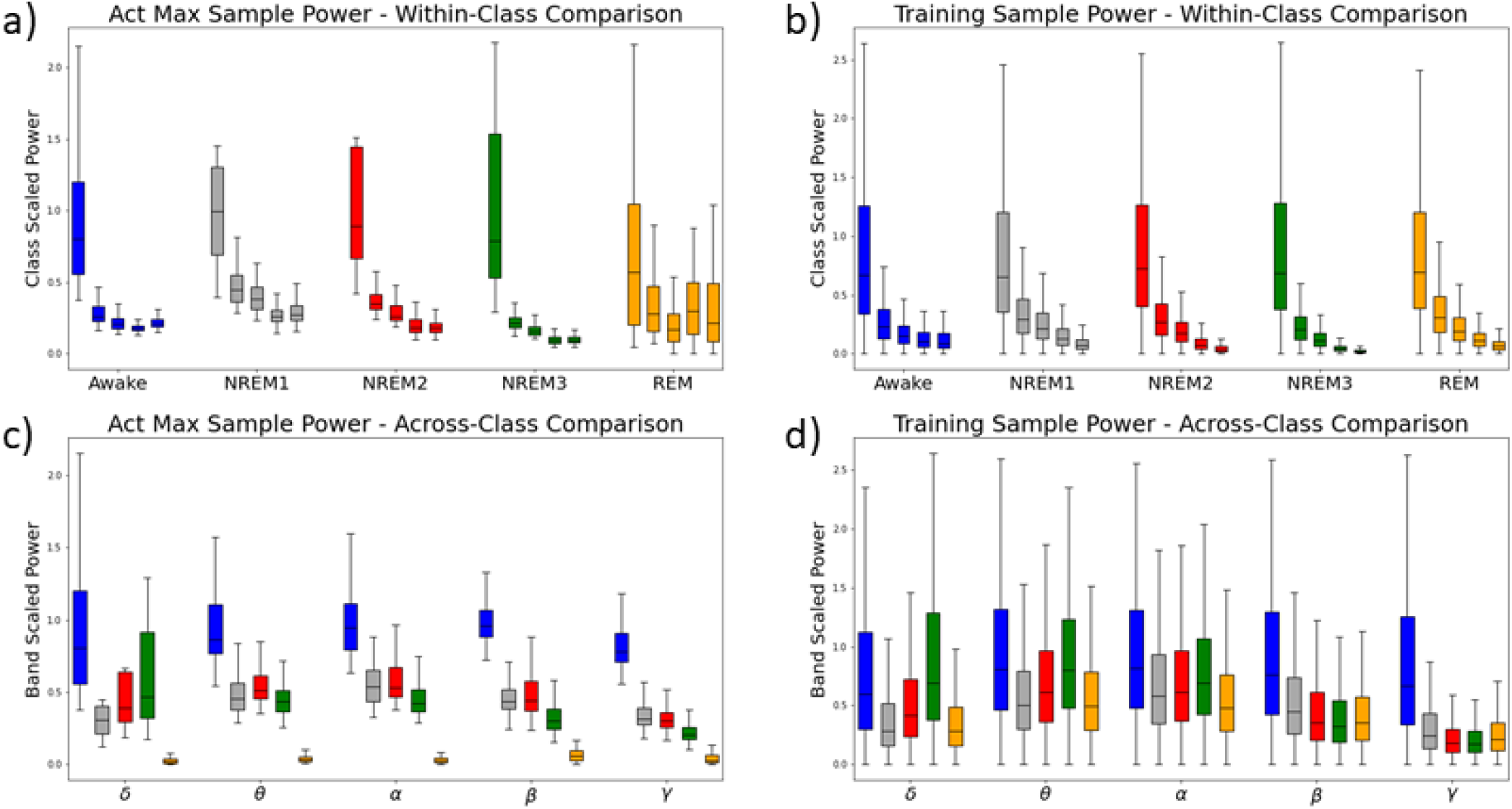
Power for Training Samples and Samples Generated via Activation Maximization. Panels a) and c) show the average power of the activation maximization samples in each frequency band and class. Panels b) and d) show the average power of the training samples in each frequency band and class. The values for each class in panels a) and b) are divided by the mean power of their respective δ bands to enable an easier comparison of the power of each frequency band within each class. The values for each frequency band in panels c) and d) are divided by the largest mean of the values for that frequency band across classes. This enables an easier comparison of the relative power of the frequency bands across classes. Blue, gray, red, green, and orange boxes refer to Awake, NREM1, NREM2, NREM3, and REM, respectively. In panels a) and b), the boxes are arranged based upon class, with each class having 5 boxes indicating δ, θ, α, β, and γ importance from left to right. In panels c) and d), the boxes are arranged based upon frequency band with each band having 5 boxes indicating Awake, NREM1, NREM2, NREM3, and REM from left to right.

Fig. 3c and Fig. 3d show the results for each of the activation maximization samples and for the training samples, respectively. The spectra are scaled within each frequency band by the mean of the class with the largest mean value for that band. Across both the training and activation maximization samples, Awake samples generally has the highest power. This is the case for all bands except for δ and θ in the training data. The activation maximization Awake sample has significantly larger amplitudes across all bands relative to other classes. It is possible that this nearly band-wide difference in power could explain the high performance of the classifier for Awake on the training data and the extremely high activations in Fig. 2a. Additionally, it is possible that activation maximization identified this reliance of the classifier upon high amplitude and simply continuously magnifies the amplitude to increase the activation. This large amplitude could also explain why the algorithm requires so many steps to reach its peak activation. After Awake, NREM3 has the highest δ power [25]. NREM2, followed by NREM1, has the highest θ power, which is slightly different than what we observe in the training data. Additionally, NREM1 and NREM2 have the highest α, β, and γ power. While NREM1 and NREM2 would not typically be expected to have the highest amplitude after Awake in these bands, it is not unreasonable given that the classes had values highly similar to NREM3 and REM in the training data.

It is important to emphasize that the spectral patterns our approach uncovered are highly similar to real data. Additionally, the patterns are much more similar to real data than the results of our initial attempts using time-domain-based activation maximization.

### E. Activation Maximization: Analysis of Frequency Importance

Figure 4 shows the importance of each canonical frequency band to the activation maximization sample generated for each class. Relative to the other classes, perturbation of the frequency bands of the Awake class has relatively little effect. We previously discussed how the classifier seemed to identify high levels of band-wide power corresponding to Awake, so it is possible that the perturbation of one band did not affect the activation of the sample given that the model could also rely upon the remaining high power of all of the other bands. NREM1 seems to rely most upon δ, so it is possible that the model identified the vertex sharp waves that occur in NREM1 and that can appear within δ. Following δ, θ and α are most important, and NREM1 is typically characterized by θ. For NREM2, the model seemed to rely most upon β followed by γ, which could indicate that the model identified the sleep spindles that appear in β. Importantly, the model clearly identifies NREM3 as most important for the NREM3 class, and the AASM considers δ to be a defining feature of NREM3. REM is typically characterized as multispectral, which fits our results. Additionally, β and θ are found to be particularly important by our approach, and research has found that they are particularly prominent in the frontal regions of the brain during REM [34].

**Fig. 4.**
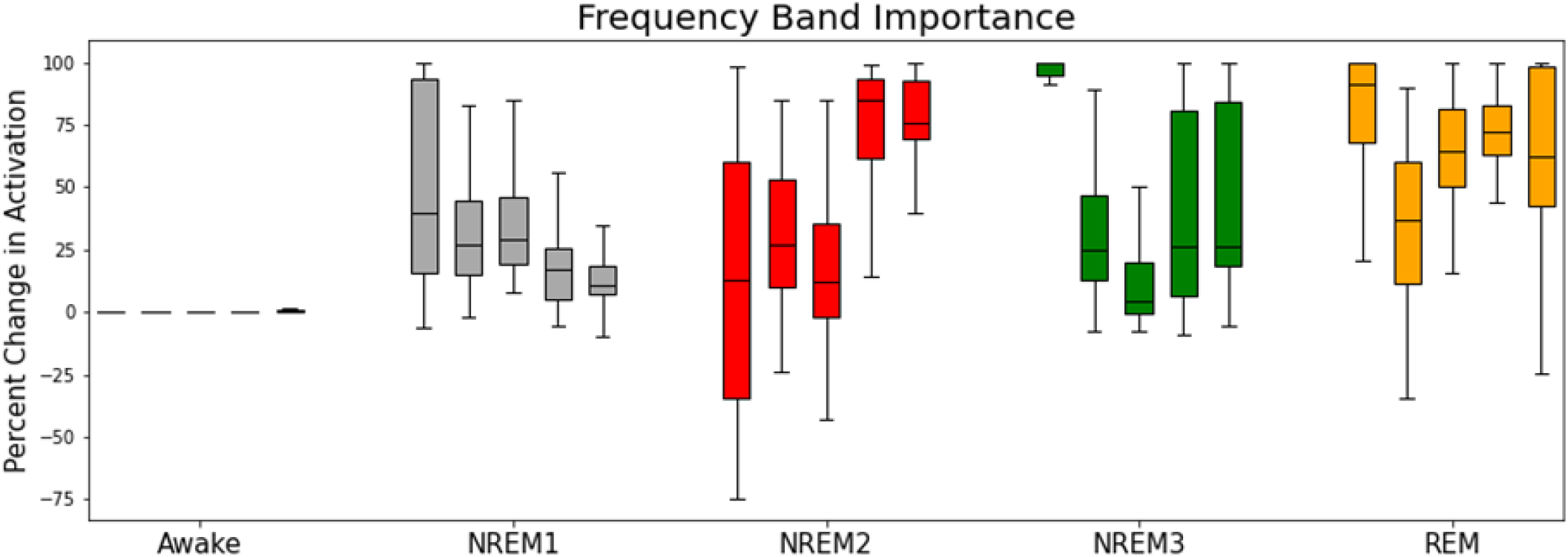
Frequency Band Importance Results for Each Class. The figure shows the percent change in activation following the perturbation of each frequency band of the generated samples. The blue, gray, red, green, and orange boxes refer to Awake, NREM1, NREM2, NREM3, and REM, respectively. The five boxes of each color refer to δ, θ, α, β, and γ importance from left to right.

### F. Further Discussion on Approach Development

The application of activation maximization to EEG time-series is novel. As such, when we first sought to develop an activation maximization approach for time-series, we optimized the time-domain content of the signal, similar to image-based analyses. However, this did not work well. When first developing the approach, we did not use boundary conditions. The use of boundary conditions enormously improved the activations. Additionally, we tested 3 initialization approaches. We initialized with the mean of each frequency, with the median of each frequency, and with zeros. Using the median of each frequency seemed to work best. Additionally, when first implementing gradient descent, we multiplied both the direction and magnitude of the gradient by the step size for each step. However, this led to results that were highly inconsistent across parameter settings. Just multiplying the step size by the direction of the gradient seemed to provide more consistent results.

### G. Broader Implications of Approach

In this study, we develop, to the best of our knowledge, the first implementation of activation maximization for raw time-series classification in EEG. Additionally, our approach is highly useful for insight into long time-series. Our approach yields significantly better results than our attempts at activation maximization in the time-domain. We evaluate our approach within the context of sleep stage classification, which has key spectral features. However, our approach should be broadly applicable to any analysis involving classification of time-series. Additionally, because our approach optimizes the spectral content of the signal, it has strong potential for domains that value the frequency characteristics of waveforms. This is particularly the case for electrophysiology analysis with modalities like local field potentials, magnetoencephalography, and EEG [4][8].

### H. Limitations and Next Steps

A key limitation of our current approach is that we only demonstrate its viability for single-channel time-series. While some domains, like sleep stage classification, typically only involve one channel, electrophysiology datasets are often multi-channel. Further development and testing will be needed to extend the method to multi-channel time-series. Additionally, we optimized the spectral content through iteratively updating the real and imaginary coefficients of the FFT. It is unclear whether optimizing phase and amplitude could yield improved results, but that is a possibility worth investigating in the future. Future efforts could adapt our approach to other domains that involve time-series analysis but that do not prioritize spectral features to the same degree.

## IV. Conclusion

Many electrophysiology analyses rely upon insight into the frequency domain. In this study, we present FAME, a novel Frequency-based Activation Maximization Explainability approach that generates samples with features characteristic of the patterns learned by a deep learning classifier for each sample. It is, to the best of our knowledge, the first adaptation of activation maximization for domains with long time-series and strong dynamics. Moreover, because it involves the iterative updating of FFT coefficients to generate samples, it provides insight into the frequencies most important to creating the samples, which is particularly important to the analysis of electrophysiology data. We demonstrate the utility of FAME within the context of sleep stage classification, and we find that it uncovers patterns known to be associated with EEG sleep stages. FAME also uncovers some patterns that may be associated with artifact and have resulted in improved performance for the Awake class. Through its creative use of frequency domain optimization, our approach could lead to the development of a new class of time-series explainability approaches. Moreover, our novel approach has the potential to greatly impact the domain of electrophysiology analysis and other fields involving long and highly dynamic time-series.

